# Superiority of intranasal over systemic administration of bioengineered soluble ACE2 for survival and brain protection against SARS-CoV-2 infection

**DOI:** 10.1101/2022.12.05.519032

**Authors:** Luise Hassler, Jan Wysocki, Jared T. Ahrendsen, Minghao Ye, Ian Gelarden, Vlad Nicolaescu, Anastasia Tomatsidou, Haley Gula, Cosimo Cianfarini, Nigar Khurram, Yashpal Kanwar, Benjamin D. Singer, Glenn Randall, Dominique Missiakas, Jack Henkin, Daniel Batlle

## Abstract

The present study was designed to investigate the effects of a soluble ACE2 protein termed ACE2 618-DDC-ABD, bioengineered to have long duration of action and high binding affinity to SARS-CoV-2, when administered either intranasally (IN) or intraperitoneally (IP) and before or after SARS-CoV-2 inoculation.

K18hACE2 mice permissive for SARS-CoV-2 infection were inoculated with 2×10^4^ PFU wildtype SARS-CoV-2. In one protocol, ACE2 618-DDC-ABD was given either IN or IP, pre- and post-viral inoculation. In a second protocol, ACE2 618-DDC-ABD was given either IN, IP or IN+IP but only post-viral inoculation. In addition, A549 and Vero E6 cells were used to test neutralization of SARS-CoV-2 variants by ACE2 618-DDC-ABD at different concentrations.

Survival by day 5 was 0% in infected untreated mice, and 40% in mice from the ACE2 618-DDC-ABD IP-pre treated group. By contrast, in the IN-pre group survival was 90%, histopathology of brain and kidney was essentially normal and markedly improved in the lungs. When ACE2 618-DDC-ABD was administered only post viral inoculation, survival was 30% in the IN+IP group, 20% in the IN and 0% in the IP group. Brain SARS-CoV-2 titers were high in all groups except for the IN-pre group where titers were undetectable in all mice. In cells permissive for SARS-CoV-2 infection, ACE2 618-DDC-ABD neutralized wildtype SARS-CoV-2 at high concentrations, whereas much lower concentrations neutralized omicron BA. 1.

We conclude that ACE2 618-DDC-ABD provides much better survival and organ protection when administered intranasally than when given systemically or after viral inoculation and that lowering brain titers is a critical determinant of survival and organ protection.

## INTRODUCTION

Early in 2020, shortly after ACE2 was reported to be the main cell entry receptor for SARS-CoV-2 (1, 2), our lab proposed the use of soluble ACE2 proteins to neutralize SARS-CoV-2 via a decoy effect (3). The potential of soluble ACE2 proteins to neutralize SARS-CoV-2 was soon after shown using human organoids (4). This cellular model expresses human ACE2, the essential cell entry receptor for SARS-CoV-2 and TMPRSS2, a protease critical for internalization of the ACE2-SARS-CoV-2 complex (2, 4–6). Since mice and rats are resistant to wildtype SARS-CoV-2, the human transgenic k18hACE2 mouse has been used widely to test the efficacy of new interventions geared to prevent and treat SARS-CoV-2 infection (7–14). The k18hACE2 model is lethal when infected with a high dose of wildtype SARS-CoV-2 and replicates severe human lung disease in humans (10, 11, 14–16). There is also some evidence of brain injury (9, 10, 14, 17–19), but the precise cause of the universal lethality is not known.

Soluble ACE2 proteins for SARS-CoV-2 offer theoretical advantages over antibody-based approaches which are increasingly resistant to emerging SARS-CoV-2 variants (20–30). For instance, multiple passaging in the presence of soluble ACE2 proteins does not lead to mutational escape of SARS-CoV-2, whereas mutational escape of the virus is seen rapidly after passaging in the presence of monoclonal antibodies (31). ACE2 decoys have a unique advantage over monoclonal antibodies, because viral mutants are unlikely to decrease decoy affinity without simultaneous loss of ACE2 affinity, making decoys less susceptible to resistance by viral mutation (32–34).

We bioengineered a soluble ACE2 protein, based on a truncate of human ACE2 with 618 amino acids that was fused with an albumin binding domain (ABD) to confer prolonged *in vivo* duration of action via albumin binding (5). Later, we used a dodecapeptide (DDC) motif to form a dimer and enhance the binding affinity for SARS-CoV-2 (8). In the k18hACE2 model infected with SARS-CoV-2, this protein, termed ACE2 618-DDC-ABD, resulted in markedly improved survival and greatly reduced lung injury (8). ACE2 618-DDC-ABD in this previous study was administered combined intranasally (IN) and intraperitoneally (IP) to ensure proof-of-concept efficacy (8). Here, we compared the IN versus IP administration of ACE2 618-DDC-ABD and examined the impact of treatment post viral inoculation on survival, organ protection and viral titers.

## RESULTS

### Survival, clinical score and weight loss in SARS-CoV-2 infected k18hACE2 mice

The effects of intranasal (IN) versus intraperitoneal (IP) administration of the soluble ACE2 protein (termed ACE2 618-DDC-ABD) were examined in the k18hACE2 mouse, a lethal model of SARS-CoV-2 infection. According to study protocol, animals that lost more than 20% of their body weight or had a clinical score of 3 or higher were humanely euthanized and this was considered a mortality event (7, 8). Suvival was 0% in the infected untreated control mice, all of which had to be humanely euthanized on day 5 (**fig 1A**), had severe body weight loss (**fig 1B**) and a high clinical score (**fig 1C**).

**Figure 1.**
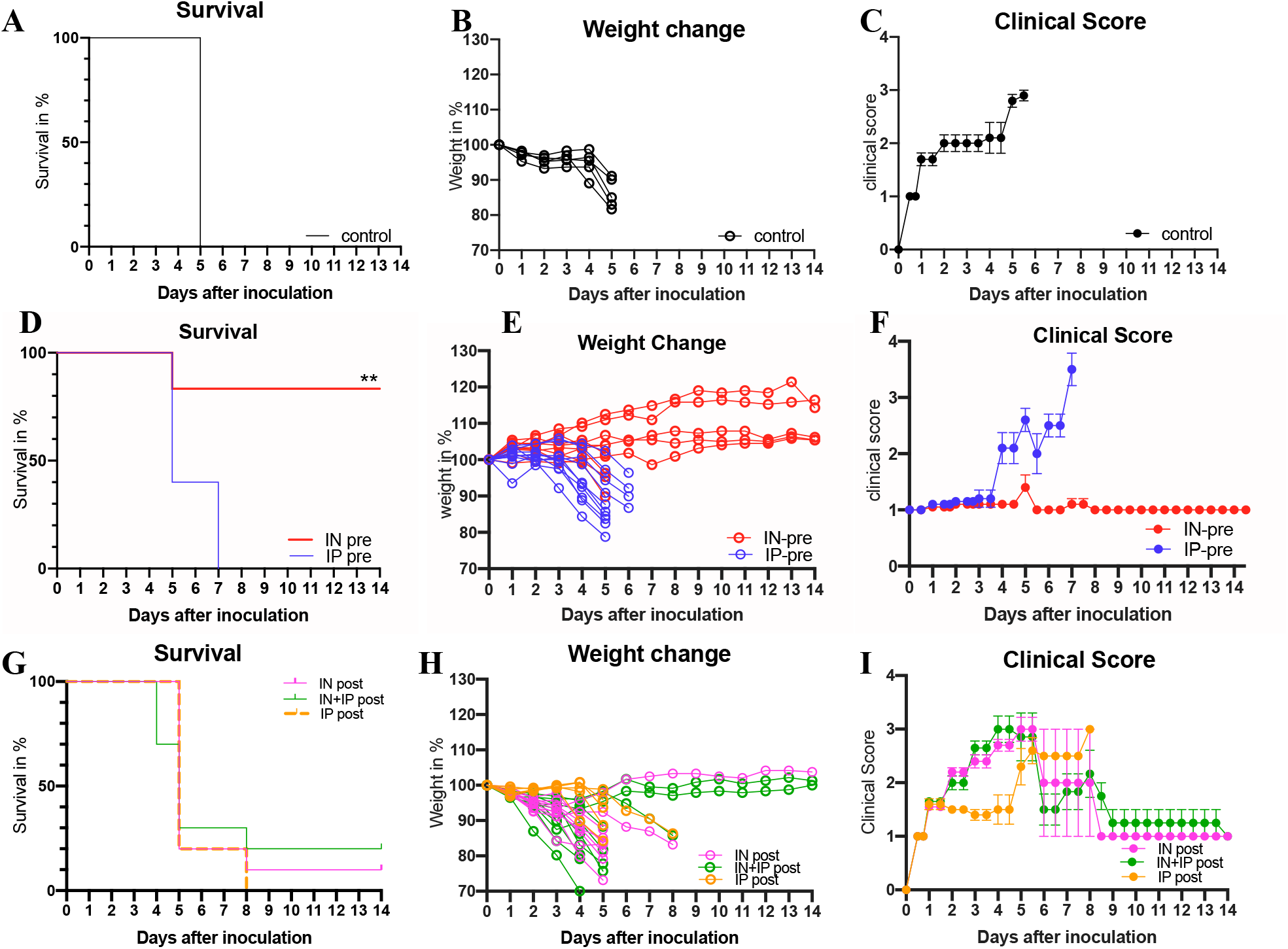
Survival, body weight and clinical score comparing intranasal (IN) vs intraperitoneal (IP) vs IN+IP ACE2 618-DDC-ABD administration to k18hACE2 mice prior to or post viral inoculation with SARS-CoV-2. **Panel A-C, vehicle treated group.** Infected mice that received vehicle (BSA in PBS, black) had 0% survival on day 5 (**A**), lost up to 20% of their body weight (**B**) and developed high clinical scores (**C**). **Panel D-F, administration prior to and post viral inoculation.** In the IN-pre group (red) nine out of ten mice survived until day five (90%), whereas in the IP-pre group (blue) only four out of ten mice survived until day five (40%). Four of the nine surviving mice from the IN-pre group that were healthy by clinical score were then sacrificed to obtain organs for comparison and the remaining five mice all survived until day 14. By contrast, none of the four remaining mice in the IP-pre group survived until day 14 (p=0.0024 by Log-rank (Mantel-Cox) test) (**D**). The IN-pre group (red) had no body weight loss (**E**) and clinical score was normal (**F**) whereas the IP-pre group (blue) experienced body weight loss and worsening clinical score. **Panel G-I, administration post viral inoculation.** In the IN+IP-post group (green), 3 out of 10 mice survived until 5 (30%) and 2 out of 10 (20%) until day 14. In the IN-post group (pink), survival was 2 out of 10 (20%) on day 5 and 1 out of 10 (10%) on day 14. In the IP-post group (orange), survival was 1 out of 5 (20%) on day 5 and 0 out of 5 (0%) on day 14 (p=0.9250 by Log-rank (Mantel-Cox) test) (**G**). In most mice that received ACE2 618-DDC-ABD post viral inoculation weight loss was severe and clinical scores were high although a few mice had stable weight and normal clinical score (**H, I**).

#### a) Administration of ACE2 618-DDC-ABD pre- and post-viral inoculation

In mice that received ACE2 618-DDC-ABD pre- and post-viral inoculation, survival on day 5 was 90% in the IN-pre group (9 out of 10), whereas it was only 40% in the IP-pre group (4 out of 10) (**fig 1D**). This is in contrast with untreated infected controls that had 0% survival on day 5 (**fig 1A**). Four of the nine mice from the IN-pre group that were healthy based on their clinical score were sacrificed on day 5 to obtain organs for comparison to the IP-pre group, and the remaining five mice from the IN-pre group all survived until day the end of the study, day 14. The remaining mice in the IP-pre-group, by contrast, all had to be humanely euthanized by day 7 because of worsening clinical scores and weight loss, as per protocol (**fig 1D**). The IN-pre-group, moreover, had no body weight loss (**fig 1E**) and better clinical scores (**fig 1F**). The IP-pre-group, by contrast, experienced body weight loss (**fig 1E**) and had worsening clinical scores (**fig 1F**).

#### b) Administration of ACE2 618-DDC-ABD post-viral inoculation

In mice that received ACE2 618-DDC-ABD only post viral inoculation, survival in the IN+IP-post group was 30% on day 5 (3 out of 10) and 20% on day 14 (2 out of 10) (**fig 1G**). In the IN-post group, survival was 20% on day 5 (2 out of 10) and 10% on day 14 (1 out of 10). In the IP-post-group, survival was also 20% on day 5 (1 out of 5) but 0% on day 14 (0 out of 5) (**fig 1G**). For comparison, infected untreated mice had 0% survival on day 5 (**fig 1A**). Most of the animals that received ACE2 618-DDC-ABD post viral inoculation had rapid body weight loss and a worsening clinical score, but some (n=2 IN+IP-post, n=1 IN-post) recovered over the course of the study and survived until day 14 (**fig 1H, I**).

### SARS-CoV-2 titers in lungs, brains and kidneys

#### a) Administration of ACE2 618-DDC-ABD pre- and post-viral inoculation

Brain titers were non-detectable in the IN-pre group (**fig 2A**). By contrast, in the IP-pre group, brain titers were high in 5 out of 8 mice, and as a group significantly higher than in the IN-pre group (3.82E+08 and 0 PFU/ml, respectively) (**fig 2A**).

**Figure 2.**
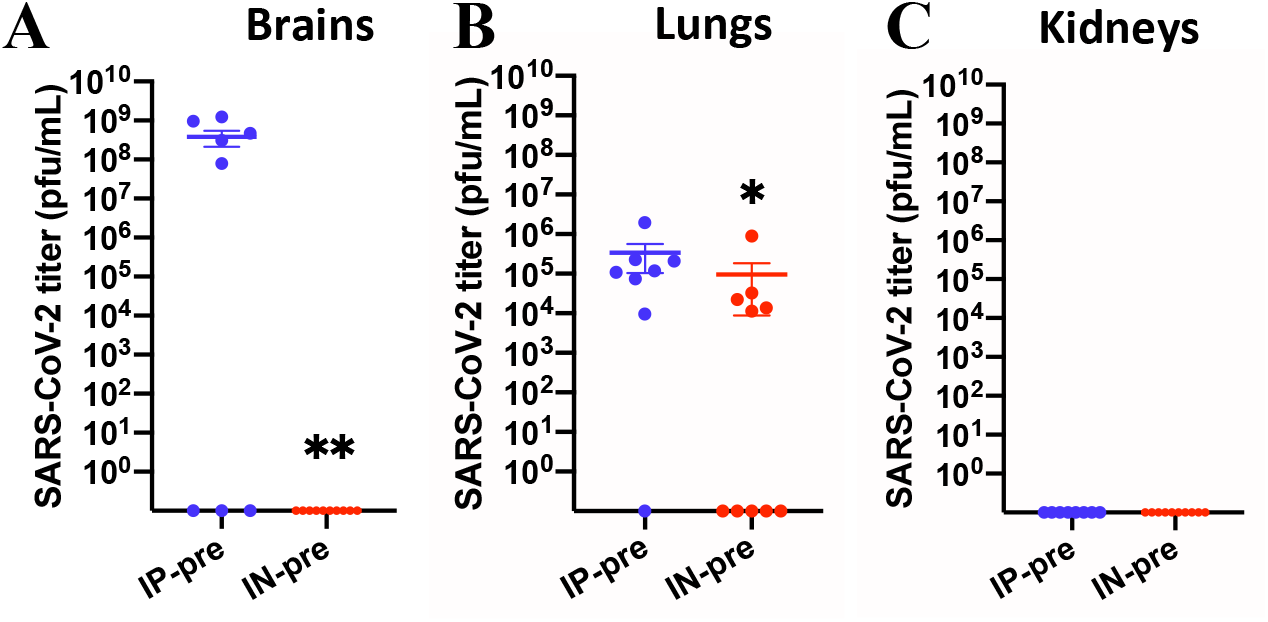
Brain, lung and kidney viral titers in SARS-CoV-2 inoculated k18hACE2 mice that received ACE2 618-DDC-ABD pre- and post viral inoculation. **Panel A.** Brain titers are high in 5 out of 8 animals of the IP-pre group (blue) whereas they are undetectable in the 10 IN-pre-treated animals (red) (p=0.0065). **Panel B.** Lung titers are slightly but significantly decreased in the IN-pre group (n=10, red) as compared to the IP-pre group (n=8, blue) (p=0.0426). **Panel C.** Kidney viral titers were non-detectable in both groups. Note that in the IP-pre group organs could be obtained in 8 out of 10 mice only. Significance was calculated using Mann-Whitney-Test for non-normally distributed data. Viral titers were normalized by organ weight. Mean ± SEM are shown.

Lung titers were significantly lower in the IN-pre group as compared to the IP-pre group (9.67E+04 and 3.37E+05 PFU/ml, respectively) (**fig 2B**).

Kidney titers were not detected in any group and at any time point (**fig 2C**).

#### b) Administration of ACE2 618-DDC-ABD post-viral inoculation

Brain viral titers were decreased only marginally and not significantly in post-treated mice as compared to infected untreated mice (**fig 3A**). In the few survivors from the post-treated groups (n=2 IN+IP-post, n=1 IN-post), however, brain titers were undetectable on day 14 (**fig 3A**).

**Figure 3.**
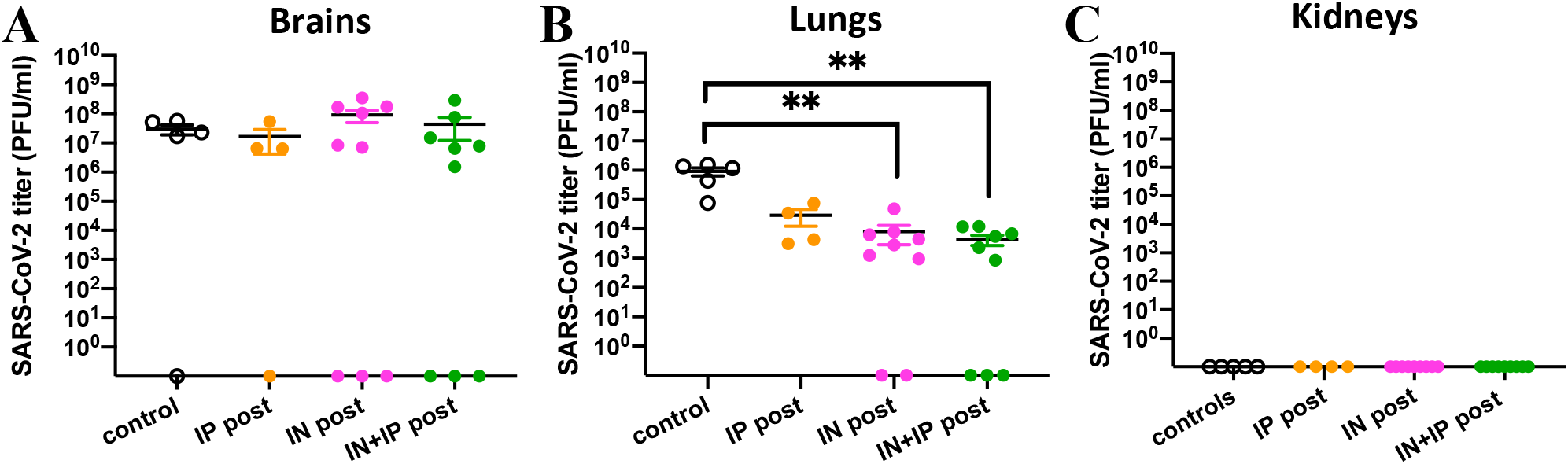
Brain, lung and kidney viral titers in SARS-CoV-2 inoculated k18hACE2 mice that received ACE2 618-DDC-ABD only post viral inoculation. **Panel A.** Brain titers were high in 4 out of 5 control animals (black). Brain titers were high in 6 out of 9 IN-post (pink) and 6 out of 9 IN+IP-post (green) treated mice. In the IP-post group (orange) titers were high in 3 out of 4 animals. Organs could not be obtained from one mouse in each treated group. There were no statistically significant differences between the groups. **Panel B.** Lung titers were high in untreated controls (black). In the IN-post (pink) and IN+IP-post treated groups (green), lung titers were significantly reduced (p=0.007 and p=0.004, respectively), whereas in the IP-post group (orange) the difference was not significant. Significance was calculated by one-way ANOVA followed by Dunn’s multiple comparisons test. **Panel C.** Kidney viral titers were non-detectable in the SARS-CoV-2 infected, untreated controls (black) as well as in the mice that received ACE2 618-DDC-ABD post-viral inoculation (color). Note that organs from one mouse per post-treated groups could not be obtained. Viral titers were normalized by organ weight in all mice. Mean ± SEM are shown.

Lung SARS-CoV-2 titers in mice treated post viral inoculation were high but significantly lower than in infected untreated mice that did not receive ACE2 618-DDC-ABD. The difference reached statistical significance for the IN-post and IN+IP-post group as compared to infected untreated controls (**fig 3B**). In the IP-post group the difference was not statistically significant as compared to infected untreated controls (**fig 3B**).

Kidney viral titers were not detected in any group and at any time point (**fig 3C**).

### Brain histopathology

In brains of the infected untreated mice the findings most consistently seen were perivascular leucocytosis and lymphocytosis and endothelial hypertrophy (**fig 4A-C**). These findings were mainly located in striatum, cortex and hypothalamus. Perivascular and parenchymal inflammation in the hypothalamus and basal ganglia was seen occasionally (**fig 4D**).

**Figure 4.**
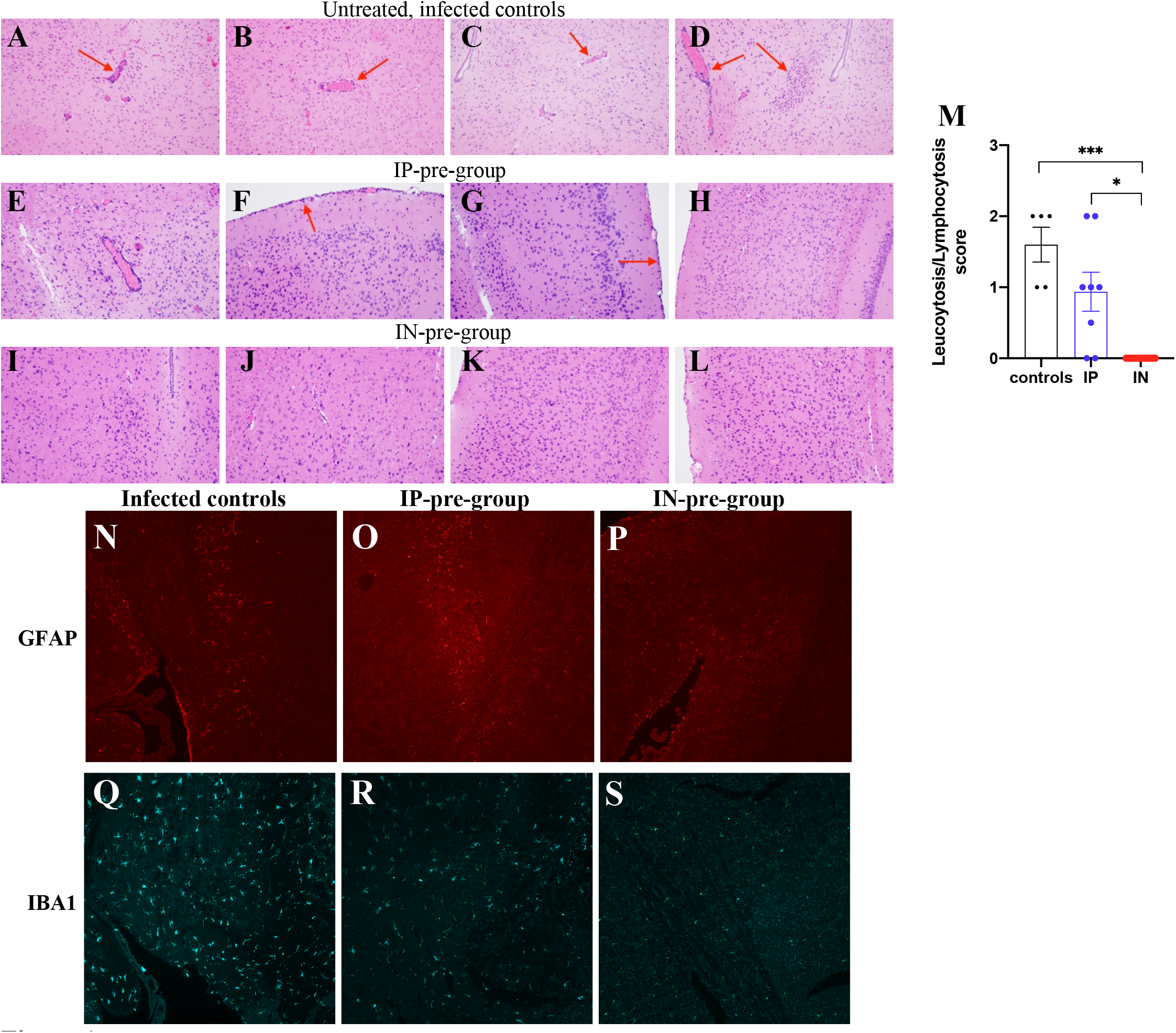
Representative neuropathology of SARS-CoV-2 infected k18hACE2 mice. **Panel A-D.** Examples of four infected untreated mice showing perivascular lymphocytosis in the ventral striatum (**A**) and lateral cerebral cortex (**B**), endothelial hypertrophy in the hypothalamus (**C**) and perivascular and parenchymal inflammation in the hypothalamus and basal ganglia (**D**). **Panel E-H.** Examples of four animals from the IP-pre group that received ACE2 618-DDC-ABD also showing perivascular lymphocytosis in the brainstem (**E**) and mild leptomeningeal lymphocytosis in the posterior (**F**) and lateral (**G**) cerebral cortex. A different mouse of the same group did not show these lesions (**H**). **Panel I-L.** Examples of four animals from the IN-pre group that received ACE2 618-DDC-ABD showing normal appearing hypothalamus (**I**), thalamus (**J**), and cerebral cortex (**K, L**). **Panel M.** When data from all animals (n=5 controls, n=8 IP-pre, n=10 IN-pre) was scored for leucocytosis/ lymphocytosis, there were significant differences between the groups (p=0.0007 and p=0.01, respectively). Mean ± SEM are shown. Significance was calculated by one-way ANOVA followed by Dunn’s multiple comparisons test. **Panel N-P.** Examples of GFAP staining (red) which is strongest in an untreated infected brain (**N**), reduced partially in a brain from the IP-pre group (**O**), and very weak in a brain from the IN-pre group (**P**). **Panel Q-S.** Examples of IBA1 staining (blue) which is strongest in an untreated infected brain (**Q**), reduced partially in a brain from the IP-pre group (**R**), and almost completely absent in a brain from the IN-pre group (**S**). All photomicrographs were taken at 20x magnification, please magnify to see the differences better.

#### a) ACE2 618-DDC-ABD administration pre- and post-viral inoculation

In some mice in the IP-pre group there was also perivascular lymphocytosis (**fig 4E**) or mild leptomeningeal lymphocytosis in the lateral and posterior cortex (**fig 4F, G**). These lesions, however, were not seen in all animals in this group (**fig 4H**). In the IN-pre group, by contrast, hypothalamus and cortex appeared normal in all mice (**fig I-L**). The differences reached statistical significance for the IN-pre group compared to both infected untreated mice and the IP-pre group (**fig 4M**).

Another abnormality found in untreated infected brains was neuronal pyknosis (examples not shown). This was scored on a scale of 0-3 and was significanlty reduced in both the IP-pre (0.875±0.125) and IN-pre group (1±0) as compared to untreated infected mice (2±0, p=0.0001 and p=0.0003, respectively).

#### Immunofluorescence for brain injury markers

Immunofluorescence for the astrocyte activation marker GFAP and the microglial activation marker IBA1 revealed expression of both markers in infected untreated mice (**fig 4N, Q**). GFAP and IBA1 staining was partially decreased in mice in the IP-pre group (**fig 4O, R**), whereas it was markedly reduced in the IN-pre group (**fig 4P, S**). Several other examples showing these differences are shown in the supplement (**fig S1**).

#### b) ACE2 618-DDC-ABD administration post-viral inoculation

In the IP-post group, perivascular and parenchymal inflammation and endothelial hypertrophy in the hypothalamus was also seen (**fig S2A, B**). In both the IN-post and IN+IP-post group brain histopathology was improved in some animals, whereas leptomeningeal and perivascular lymphocytosis and endothelial hypertrophy in hypothalamus and cortex were seen in others similar to the infected untreated mice (**fig S2C-F**).

The score for leucocytosis/lymphocytosis was lower in the IP-post, IN-post and IN+IP-post group than in the infected untreated mice (**fig S2G**). These differences, however, did not reach statistical significance (**fig S2G**). The score for neuronal pyknosis was decreased in all three treated groups, but the difference was significant only for the IN+IP-post group as compared to infected untreated mice (**fig S2H**).

### Lung histopathology

Lungs from untreated infected mice showed dense perivascular mononuclear infiltrates and collections of intra-alveolar neutrophils. There were also rare foci of necrotic debris and alveolar hemorrhage (**fig 5A-E**).

**Figure 5.**
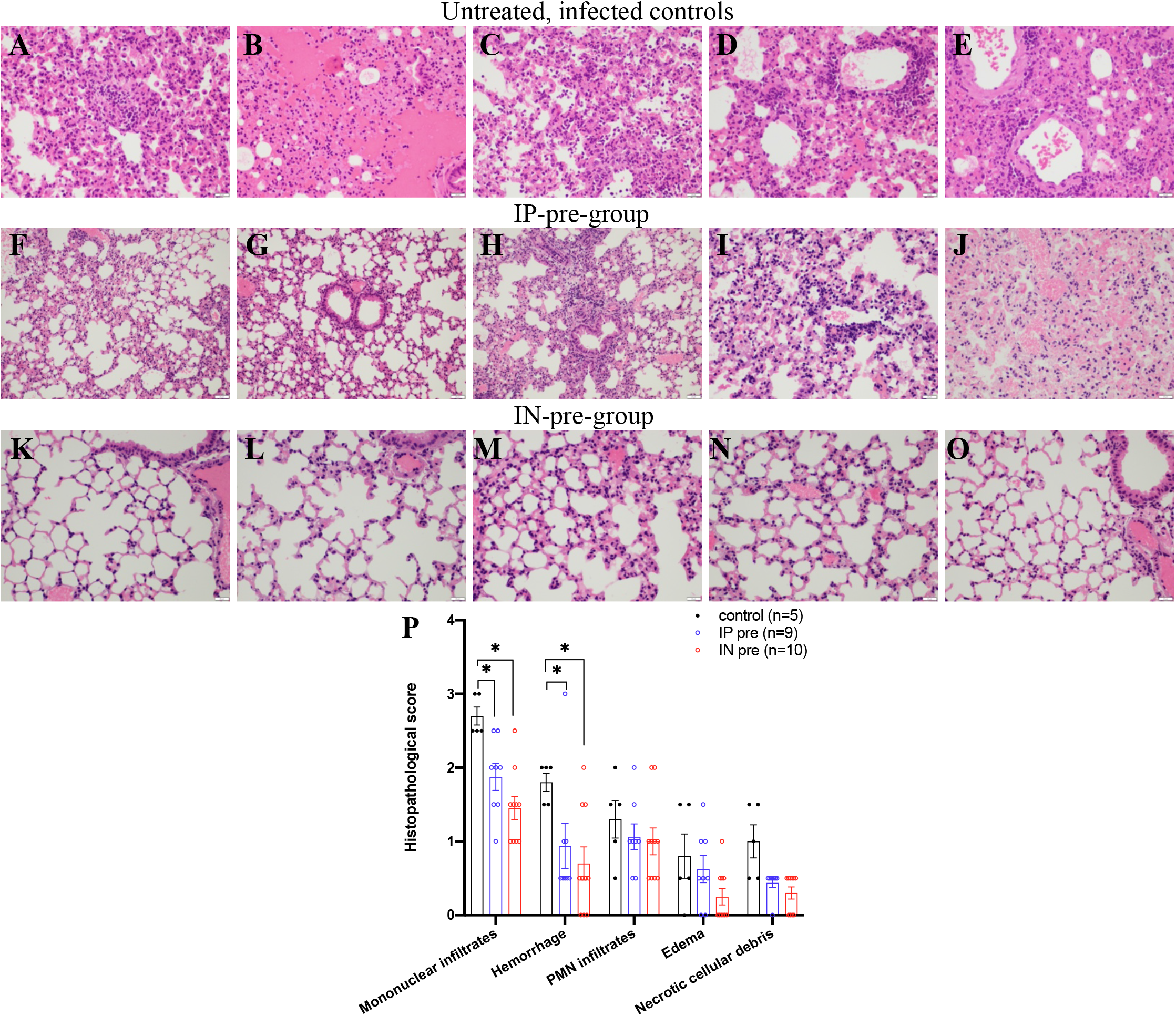
Representative examples of lung histopathology in SARS-CoV-2 infected k18hACE2 mice. **Panel A-E.** Five different mice from the **vehicle-control group** (n=5) that were infected with SARS-CoV-2 show dense perivascular mononuclear infiltrates (**D, E**), collections of intra-alveolar neutrophils (**C**), rare foci of necrotic debris (**A**) and alveolar hemorrhage (**B**). **Panel F-J.** Five different mice from the group that received **ACE2 618-DDC-ABD pre-and post viral inoculation via IP** (n=8) show milder perivascular mononuclear infiltrates (**H, I**) with other areas of near normal lung histopathology (**F, G**), focal minimal alveolar hemorrhage (**J**), and occasional intra-alveolar neutrophils. **Panel K-O.** Five different mice from the group that received **ACE2 618-DDC-ABD pre- and post viral inoculation via IN** (n=10) show near normal lung histopathology with minimal (**K-O**) perivascular mononuclear infiltrates and hemorrhage in some cases. **Panel P.** The lung histopathology scores for mononuclear infiltrates, hemorrhage, PMN infiltrates, edema and necrotic cellular debris are high in the controls (black) and lower in both the IP-pre (blue) and IN-pre (red) group. Mean ± SEM are shown. Significance is indicated in the figure with * = p<0.05, calculated by mixed-effects analysis followed by Tukey’s multiple comparisons test. All photomicrographs were taken from H&E-stained sections at 40x magnification, scale bar = 500um.

#### a) Administration of ACE2 618-DDC-ABD pre- and post-viral inoculation

The lungs from mice from the IP-pre group showed few perivascular mononuclear infiltrates, focal minimal alveolar hemorrhage and occasional intra-alveolar neutrophils, whereas some areas also show near normal lung histopathology (**fig 5F-J**). The lungs from mice from the IN-pre group show near normal lung histopathology with only minimal perivascular mononuclear infiltrates (**fig 5K-O**). The lung histopathology scores for mononuclear infiltrates, hemorrhage, PMN infiltrates, edema and necrotic cellular debris were worse in the untreated infected group than in both the IP-pre and IN-pre treated groups (**fig 5**). The differences in both pre-treated groups were highly significant for the main histopathologic findings mononuclear infiltrates and alveolar hemorrhage, as compared to infected untreated mice (**fig 5P**). The scores were better in the IN-pre group than the IP-pre group, but the difference did not reach statistical significance (**fig 5P**).

#### b) Administration of ACE2 618-DDC-ABD post viral inoculation

The lungs from the post-treated groups also showed perivascular mononuclear infiltrations and intra-alveolar neutrophils and resemble the infected untreated group (**fig S3A-C**). In some mice from the IN-post and IN+IP-post group (**fig S3B, C**), lung histopathology is improved but to a lesser extent than in the pre-treated groups (**see fig 5**). The histopathological scores were not significantly different in infected untreated controls as compared to the post-treated groups (**fig S3D**). The alveolar hemorrhage was less seen in the post-treated groups, but the difference was not statistically significant (**fig S3D**).

### Kidney histopathology and markers of proximal tubular injury

Examination of the kidneys from infected untreated mice sacrificed on day 5 post viral inoculation showed mild proximal tubular injury in PAS-stained sections consisting of attenuation of proximal tubular brush border, cytolysis and tubular basement membrane disruption (**fig S4A**). These changes were variable from one mouse to the other and sometimes unremarkable. On a score of 0-3, 0 being normal, the mean score for brush border loss was 1.6±0.4 (**fig S4**).

NGAL staining, as expected, was absent in a normal uninfected kidney but strong in a kidney with AKI caused by ischemia-reperfusion injury (IRI) (30 minutes), both used here as examples for comparison (**fig S5A**). In the infected untreated mice NGAL staining was evident in all animals but variable in its intensity (**fig S5B**). KIM-1 staining was completely absent in a normal kidney and positive in a kidney with AKI caused by IRI (**fig S7A**). In infected untreated mice it was absent in all but one mice (**fig S7A, B**).

#### a) Administration of ACE2 618-DDC-ABD pre- and post-viral inoculation

In mice from both the IP-pre and IN-pre group, proximal tubular injury in PAS-stained sections was either absent or less pronounced than in infected untreated mice (**fig S4B, C**).

In the IN-pre group, NGAL staining was completely absent or very weak (**fig S5D**). In the IP-pre group, by contrast, NGAL staining was seen although was it was weak in some animals (**fig S5C**). KIM-1 staining was completely absent in both the IP-pre and IN-pre group (**fig S7C, D**).

#### b) Administration of ACE2 618-DDC-ABD post viral inoculation

Proximal tubular injury in PAS-stained sections was somewhat less pronounced than in infected untreated mice (**fig S4D-F**), but the differences were statistically significant only in some groups (**fig S4G**). NGAL staining was present in the IN-post, IP-post and IN+IP-post group, but each group had some mice in which staining was weak (**fig S6A-C**). KIM-1 staining was completely absent in all three post-treated groups (not shown). As noted above, KIM-1 staining was also negative in the infected, untreated group except for one mouse.

### ACE2 618-DDC-ABD neutralizes wildtype and omicron SARS-CoV-2 in two cell types

In human A549 cells, ACE2 618-DDC-ABD neutralized wildtype SARS-CoV-2 (as shown by cell viability) at high concentrations (40 and 200ug/ml). Lower concentrations (1.6 and 8ug/ml) neutralized infection only partially and very low concentrations (0.0128-0.32ug/ml) had no effect on cell viability (**fig 6A**).

**Figure 6.**
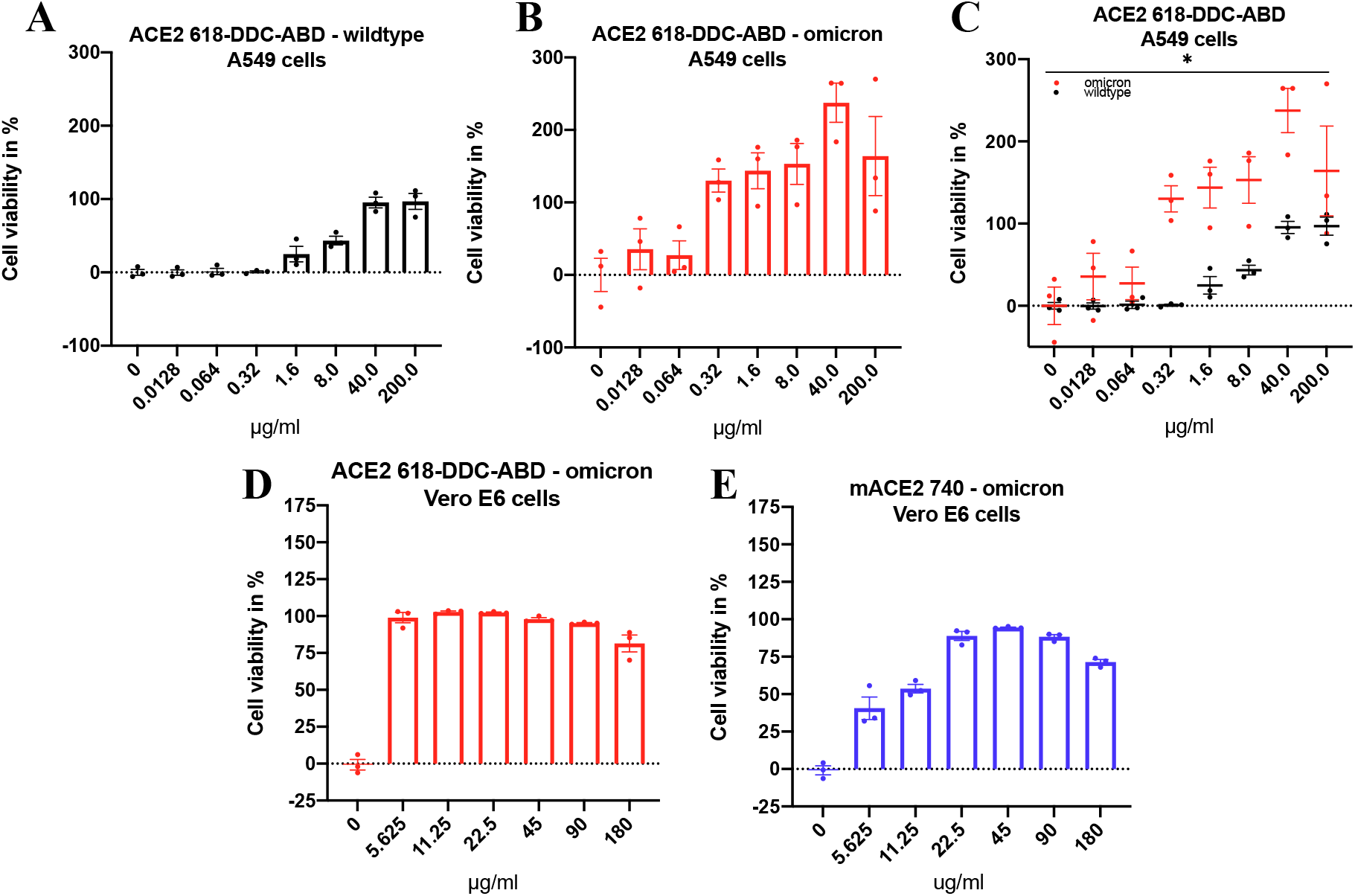
Neutralization of SARS-CoV-2 variants by human and mouse soluble ACE2 proteins. **Panel A.** In A549 cells, human ACE2 618-DDC-ABD neutralizes wildtype SARS-CoV-2 at high concentrations (40 and 200ug/ml) whereas lower concentrations (1.6 and 8ug/ml) neutralize infection only partially and very low concentrations (0.0128-0.32ug/ml) have no effect on infectivity. **Panel B.** In A549 cells, human ACE2 618-DDC-ABD neutralizes the omicron BA.1 variant of SARS-CoV-2 at concentrations lower than wildtype SARS-CoV-2 (0.32-200ug/ml), and even very low concentrations have a partial effect. **Panel C.** The differences between wildtype (black) and omicron BA.1 (red) neutralization by ACE2 618-DDC-ABD are significant (p=0.0144, calculated by two-way ANOVA). **Panel D.** In Vero E6 cells, ACE2 618-DDC-ABD neutralizes the omicron BA.1 variant at all concentrations tested (5.625-180ug/ml). **Panel E.** In Vero E6 cells, mouse ACE2 740 neutralizes infection with the omicron BA.1 variant but is only fully effective at concentrations of 22.5-180ug/ml, whereas lower concentrations (5.625 and 11.25 ug/ml) have a partial effect. Mean ± SEM are shown.

By contrast, the omicron BA.1 variant was neutralized by much lower concentrations of ACE2 618-DDC-ABD than wildtype SARS-CoV-2 (**fig 6B**). When comparing the two sets of data (omicron and wildtype), the difference was highly significant (p=0.01) (**fig 6C**).

This enhanced effect of ACE2 618-DDC-ABD on the neutralization of the omicron variant in A549 cells was also found when we used Vero E6 cells, a non-human primate cell line that has been widely used for infectivity studies with SARS-CoV-2 (4, 8). In Vero E6 cells, ACE2 618-DDC-ABD neutralized the omicron BA.1 variant at all concentrations tested (5.625-180ug/ml) (**fig 6D**).

The high sensitivity of this omicron variant to ACE2 618-DDC-ABD prompted us to test a mouse soluble ACE2 protein that normally has no effect on wild type SARS-CoV-2 infectivity (5). The mouse ACE2 740 protein neutralized omicron BA.1 infection fully at high concentrations whereas lower concentrations were only partially effective (**fig 6E**).

## METHODS

### *In vivo* infectivity studies

All work with live SARS-CoV-2 in k18hACE2 mice was performed in the BSL-3 facility of the Ricketts Regional Biocontainment Laboratory, following a protocol approved by the Institutional Animal Care and Use Committees of Northwestern University and University of Chicago. We used k18-hACE2 mice that express full length human ACE2 and are susceptible to SARS-CoV-2 infection (10, 14–16, 35, 36), purchased from Jackson Lab (8-13 weeks old). Animals were infected with 2×10^4^ PFU SARS-CoV-2 (novel coronavirus (nCoV)/Washington/1/2020 was provided by N. Thornburg (CDC) via the World Reference Center for Emerging Viruses and Arboviruses (Galveston, TX)) in 20μl by intranasal inoculation. Animals infected with this viral load invariably succumb to disease by days 5-9 (10, 14–16). We used different protocols to examine pre-treatment and post-treatment effects of the soluble ACE2 618-DDC-ABD protein and to compare intranasal (IN) versus intraperitoneal (IP) administration effects. In the pre-treatment groups, ACE2 618-DDC-ABD was administered to k18hACE2 mice (n=10, 5 male and 5 female) via IN (30μl, ~13μg/g BW) or via IP (200μl, ~13μg/g BW) 1 hour prior to SARS-CoV-2 followed by the same dose 24 and 48 hours later for a total of 3 doses. In the post-treatment groups, ACE2 618-DDC-ABD was administered either IN (30μl, ~12μg/g BW, n=10) or IP (200μl, ~1μg/g BW, n=5) or combined intranasally and intraperitoneally (IN+IP) (n=10) 24, 48 and 72 hours only post viral inoculation (2×10^4^ PFU SARS-CoV-2). Controls (n=5) received bovine serum albumin (BSA) in PBS both IN and IP at the same doses and time-points as the ACE2 618-DDC-ABD post-treated animals.

Animals were weighed once daily and monitored twice daily for health using a clinical scoring system (Table 1, supplement). Animals that lost more than 20% of their baseline body weight or had a clinical score of 3 were sacrificed for humane reasons and considered a fatal event as per study protocol. To be able to compare viral titers and organ pathology at the same time point, randomly selected animals from the IN group (which all appered healthy based on clinical score) were sacrificed on day 5 together with the animals from the IP-group that were sacrificed because the mortality endpoint was reached. Otherwise, animals that did not reach the severity of these criteria were monitored for up to 14 days in the BSL-3 facility and sacrificed at day 14.

Portions of lungs, kidneys and brains were removed from all sacrificed animals and were used for viral load measurements by plaque assay (see below), whereas the remaining portions were fixed in 10% formalin and embedded in paraffin for histopathology and immunostaining. The Mouse Histology & Phenotyping Laboratory center at Northwestern University generated slides for staining studies.

Hematoxylin and eosin (H&E)-stained sections were evaluated by expert lung pathologists on a scoring system recently described for SARS-CoV-2 infected k18-hACE2 mice (8, 14). The categories scored were: mononuclear infiltrates, alveolar hemorrhage, edema, cellular necrosis, hyaline membranes and thrombosis. The scale was as follows: 0 = no detection, 1 = uncommon detection in <5% lung fields (200x), 2 = detectable in up to 30% of lung fields, 3 = detectable in 33-66% of lung fields and 4 = detectable in >66% of lung fields. Neutrophil infiltration was evaluated on a scale of 0-3 as follows: 0 = within normal range, 1 = scattered PMNs sequestered in septa, 2 = score 1 and solitary PMNs extravasated in airspaces, 3 = score 2 plus and aggregates in vessel and airspaces.

Brain injury was evaluated on H&E-stained sections by a blinded neuropathologist and scored for leucocytosis and lymphocytosis as well as neuronal pyknosis on a scale from 0-3. The scale was as follows: 0 = none, 1 = mild (focal), 2 = moderate (multifocal), 3 = severe (diffuse). Kidney histopathology was evaluated on periodic acid schiff (PAS)-stained sections by an expert kidney pathologist for brush border loss, cytolysis and tubular basement membrane (TBM) attenuation and using two markers of proximal tubule injury, KIM-1 and NGAL (see immunofluorscence section below).

### Plaque assay for infectious virus

Tissue samples were collected in DMEM with 2% FBS and were homogenized using 1.4mm ceramic beads in a tissue homogenizer using two 30s pulses. Samples were then centrifuged at 1000g for 5 minutes and the supernatant was serially diluted 10-fold and used to infect VeroE6 cells for 1 hour. Inoculum was removed and 1.25% methylcellulose DMEM was added to the cells and incubated for 3 days. Plates were fixed in 1:10 formalin for 1 hour and stained with crystal violet for 1 hour and counted to determine plaque forming units (PFU) and expressed as PFU/ml after the data was normalized by organ weight.

### Immunofluorescence

For immunofluorescence staining studies of the brain, an IBA1 (Abcam, ab178846) and GFAP (Abcam, ab4674) antibody were used. For staining studies of the kidney, an Anti Lipocain-2/NGAL antibody (Abcam, ab216462) and a KIM-1 antibody (R&D systems AF1817, a gift from Dr. Joseph Bonventre) were used. One example of a healthy uninfected and a mouse with acute kidney injury (AKI) 48 hours after 30 minutes of ischemia reperfusion injury are included in the KIM-1 and NGAL stainings for comparison.

### SARS-CoV-2 infection of A549 and Vero E6 cells

All work with live SARS-CoV-2 was performed in hACE2-A549 or Vero E6 cells in the BSL-3 facility of the Ricketts Regional Biocontainment Laboratory. 500 PFU of each SARS-CoV-2 strain: wildtype (novel coronavirus (nCoV)/Washington/1/2020 was provided by N. Thornburg (CDC) via the World Reference Center for Emerging Viruses and Arboviruses (Galveston, TX)) or omicron BA.1 (BEI NR-56481, obtained through BEI Resources, NIAID, NIH: SARS-Related Coronavirus 2, Isolate hCoV-19/USA/GA-EHC-2811C/2021 (Lineage B.1.1.529; Omicron Variant), NR-56481, contributed by Mehul Suthar) of SARS-CoV-2 was incubated with various concentrations (0.0128, 0.064, 0.32, 1.6, 8.0, 40.0, 200ug/ml for A549 cells or 5.626, 11.25, 22.5, 45, 90, 180ug/ml for E6 cells) of the different soluble ACE2 proteins (human ACE2 618-DDC-ABD or mouse ACE2 740) for 1 hour at 37°C. This mixture was then used to infect the respective cell types. Cells were then incubated for 3-4 days (wildtype SARS-CoV-2) or 5 days (omicron BA.1) until a noticeable cytopathic effect was observed in control wells (0μg/ml of soluble ACE2 proteins). Cell numbers were assessed by staining cells with crystal violet and reading absorbance of each well at 595 nm. Values were then normalized to the 0μg/ml control and expressed as a percentage of mock (no virus) control wells.

### Statistics

GraphPad Prism v8.4.3 (GraphPad Software) was used to calculate statistics. Normality was tested using the Shapiro–Wilk test. Differences between more than two groups with normally distributed data were analyzed by ANOVA followed by post hoc Dunnett’s multiple comparisons test. Differences between more than two groups with non-normally distributed data were analyzed by the Kruskal–Wallis test followed by the post hoc Dunn’s multiple comparisons test. Differences between two groups with normally distributed data were analyzed by unpaired t-test. Differences between two groups with non-normally distributed data were analyzed by Mann-Whitney-test.

### Study approval

All work with live SARS-CoV-2 in k18hACE2 mice was performed in the BSL-3 facility of the Ricketts Regional Biocontainment Laboratory, following a protocol approved by the Institutional Animal Care and Use Committees of both Northwestern University (IS00004795) and University of Chicago (72642).

## DISCUSSION

The main finding of this study is that a soluble ACE2 protein showed clear superiority of intranasal over systemic (intraperitoneal) administration in the k18hACE2 mouse model of SARS-CoV-2 infection. This was shown by improvements in survival, clinical scores and viral titers in lungs but particularly in the brains in the groups that received the treatment intranasally. The protein, termed ACE2 618-DDC-ABD, was bioengineered to have increased duration of action by fusing a 618 amino acid truncate with an ABD-domain and designed to have increased binding affinity to the S1 spike of SARS-CoV-2 by using a DDC dimerization motif. Administration of this protein prior to viral inoculation, moreover, was far more effective regarding all the outcomes than administration only post viral inoculation.

The k18hACE2 model in which human ACE2 transgene expression is driven by the k18 promoter is lethal after 5-7 days of inoculation with a high dose of wildtype SARS-CoV-2 (8, 10, 11, 14–16). The precise cause of the rapid lethality upon SARS-CoV-2 infection in this model is not fully understood. Brain SARS-CoV-2 titers in the infected untreated control group were one order of magnitude higher than in the lungs (fig 3A compared to 3B). The absence of brain titers in mice treated intranasally explains, in our opinion, the much better survival than in untreated and groups treated with ACE2 618-DDC-ABD intraperitoneally. The group pre-treated by IN administration was indeed the only group in which brain viral titers were not detectable in any of the mice studied (fig 2A). High brain viral titers, by contrast, were detected in the other groups with the exception of a few mice that survived until day 14 and, of note, these survivors had no brain viral titers detectable. Brain viral titers, therefore, were associated with poor outcomes in terms of survival in both the pre- and post-viral inoculation groups. From these observations we conclude that improved survival conferred by ACE2 618-DDC-ABD appears to be determined by two main factors: route of administration (IN better than IP) and timing (pre-and post-viral inoculation better than only post-viral inoculation).

Consistent with our data, previous studies had suggested that brain invasion of SARS-CoV-2 in k18hACE2 mice may be associated with more severe disease (17, 18). It should be pointed out that brain injury was limited to only few animals in some studies (10, 14). Some of the findings that have been reported include encephalitis with leukocyte infiltration, hemorrhage, neuronal cell death, necrosis and spongiosis (10, 14, 17–19, 37). In these previous studies with the k18hACE2 mice (10, 17–19, 37), however, no therapies were given. Therefore, the brain findings could not be assessed regarding responses to therapies regarding improved surival and organ protection. Here, we were able to show that survival conferred by the administration of our soluble ACE2 protein was associated with non-detectable brain viral titers and better survival. Consistent with the importance of viral brain invasion, when the k18hACE2 mice were inoculated intracranially with low doses of SARS-CoV, there was lethality despite little infection in the lungs (38). When SARS-CoV-2 was administered to k18hACE2 mice in aerosolized form for more direct lung delivery and despite robust viral replication in the respiratory tract with airway obstruction, there was markedly reduced fatality and viral neuroinvasion (39). Our findings with intranasal delivery of soluble ACE2 prior to viral inoculation demonstrate the importance of obliterating brain SARS-CoV-2 invasion for survival and organ protection.

We wish to point out, however, that the brain injury was subtle even in untreated infected mice. The brain findings most frequently seen were perivascular and leptomeningeal lymphocytosis, endothelial hypertrophy and perivascular and parenchymal inflammation (fig 4). Immunofluorescence for brain injury markers in infected untreated mice revealed expression of IBA1, a marker of microglia activation, and GFAP, a marker of astrocyte activation (40, 41). Markers for astrocyte and microglia activation were also found in a study that examined cerebrospinal fluid from patients with severe Neuro-COVID (42). In studies that used immunohistochemistry and imaging mass cytometry to examine brains from deceased COVID-19 patients, astrocyte and microglia activation was found as well (43, 44). In the IN group that received the treatment prior to viral inoculation, most brains appeared normal and IBA1 and GFAP expression was decreased or absent suggesting prevention of astrocyte and microglia activation. Despite clear improvement in these parameters in mice treated with intranasal ACE2 618-DDC-ABD prior to viral inoculation it remains to be determined, however, how brain viral invasion is associated with high mortality without more evident and severe histological damage in untreated infected mice.

This study also shows that SARS-CoV-2 infected mice have variable degrees of mild proximal tubular kidney injury, namely loss of the proximal tubular brush border. The kidney injury marker NGAL was expressed in all kidneys of SARS-CoV-2 infected untreated mice consistent with our previous study in the same mouse model (8). Here we show that administration of soluble ACE2 618-DDC-ABD is associated with weaker or absent NGAL staining, mainly when given intranasally and prior to viral inoculation (fig S5D) (8). KIM-1 staining, a proximal tubular injury marker, however, was absent in most animals wether untreated or treated with ACE2 618-DDC-ABD. It has been shown that there is expression of NGAL in the lung in ventilator-associated lung injury (45). The kidney NGAL expression that we observed in untreated animals, therefore, could stem, at least in part, from extrarenal sources like the lungs (8). KIM-1 therefore may be a more specific marker of proximal tubular injury than NGAL in the setting of COVID-19. Kidney viral titers, moreover, were consistently absent in any of the infected mice in this and in a previous study (8).

The mechanisms whereby soluble ACE2 proteins can neutralize SARS-CoV-2 have been previously discussed by us and others (3, 46). ACE2 exists in two forms, a full-length membrane bound form and a shorter soluble form that lacks the transmembrane domain (47, 48) and circulates in the blood in very small amounts (49). Both forms bind the receptor binding domain (RBD) of the SARS-CoV-2 S1 spike protein. By administering an abundant amount of soluble ACE2, the spike protein of SARS-CoV-2 can be intercepted from binding to the membrane bound ACE2 by the so-called decoy effect (3). To increase the binding affinity of ACE2 618-DDC-ABD to the RBD of the SARS-CoV-2 S1 spike, a dodecapeptide (DDC) motif was introduced that induces dimerization (8). By fusion with an albumin domain binding (ABD)-tag, moreover, increased duration of action was achieved as demonstrated by its preserved plasma enzymatic activity for several days (5, 8). Membrane bound and soluble ACE2, including ACE2 618-DDC-ABD, metabolize Angiotensin II and des-Arg^9^ Bradykinin, two peptides that may be detrimental when accumulating (46, 50–52). This action may be especially beneficial in COVID-19 where internalization of ACE2-SARS-CoV-2 complexes causes depletion of cell membrane ACE2 which fosters accumulation of these pro-inflammatory peptides (52–54). Unfortunately, we were unable to measure these peptides because organ tissues could not be released from the BSL-3 facility.

The membrane bound full-length ACE2 is essential for facilitating SARS-CoV-2 infection (6, 55). As shown by previous work with the soluble human ACE2 740 (6) and here with ACE2 618-DDC-ABD in two different permissive cell types, A549 and Vero E6 cells, high concentrations of soluble ACE2 are needed to neutralize infection of cells with wildtype SARS-CoV-2. Other variants of SARS-CoV-2, however, may be effectively treated with lower doses of soluble ACE2 proteins. This can be inferred from our findings in two different permissive cell lines, where ACE2 618-DDC-ABD neutralizes the omicron BA.1 variant at lower protein concentrations (at least 20-fold lower than those required to neutralize wildtype SARS-CoV-2). It is also important to emphasize that soluble ACE2 protein-based approaches have universal effects against all the variants of SARS-CoV-2 (33). This is contrary to monoclonal antibodies that have the limitation of becoming less efficacious with each mutation of SARS-CoV-2, as consistently shown for the omicron variants (20–30). Therefore, soluble ACE2 based therapies are likely to provide universal efficacy against all SARS-CoV-2 variants that evade monoclonal antibodies.

We conclude that ACE2 618-DDC-ABD provides much better survival and organ protection when administered intranasally than when given systemically or after viral inoculation. Abrogating brain SARS-CoV-2 invasion is a critical determinant of survival and organ protection in the k18hACE2 mouse model of lethal SARS-CoV-2 infection. Treatment post-viral inoculation, while less effective, still provides improved survival and partial organ protection.

## AUTHOR CONTRIBUTIONS

L.H. participated in the design of experiments, data analysis and writing of the manuscript. J.W. designed and constructed the soluble ACE2 protein and was involved with the writing of the paper. J.A. evaluated the brain histology. M.Y. performed kidney and brain stainings. I.G. and N.K. evaluated the lung histology. G.R. supervised the infectivity studies in the transgenic mice and cells in the Biosafety Level 3 facility performed by A.T., V.N. and H.G.. D.M. provided overall supervision for the experiments in the BSL-3 facility. C.C. and Y.K. evaluated the kidney histology and immunofluorescence. B.S. was involved with study discussions and manuscript writing. J.H. provided guidance with the protein design and analysis of the findings. D.B. supervised the overall project, designed experiments and wrote most of the paper.

## ACKNOWLEDGEMENTS

L.H. and C.C. were supported by the Biomedical Education Program during a part of their stay in Chicago. Grant support to D.B. from NIH (1R21 AI166940-01) and a gift from the Joseph and Bessie Feinberg Foundation.

