## Supplement for "Superiority of intranasal over systemic administration of bioengineered soluble ACE2 for survival and brain protection against SARS-CoV-2 infection"

**Table of content:**

- **Figure S1.** Examples of GFAP and IBA1 staining in the brain of SARS-CoV-2 inoculated k18hACE2 mice.

- **Figure S2.** Representative brain histopathology of SARS-CoV-2 infected k18hACE2 mice that received ACE2 618-DDC-ABD only post viral inoculation.

- **Figure S3.** Representative lung histopathology in SARS-CoV-2 infected k18hACE2 mice that received ACE2 618-DDC-ABD only post viral inoculation.

- **Figure S4.** Representative kidney histopathology of SARS-CoV-2 infected k18hACE2 mice.

- **Figure S5.** NGAL staining in kidneys from SARS-CoV-2 infected k18hACE2 mice that received ACE2 618-DDC-ABD pre- and post-viral inoculation.

- **Figure S6.** NGAL staining in kidneys from SARS-CoV-2 infected k18hACE2 mice that received ACE2 618-DDC-ABD only post viral inoculation.

- **Figure S7.** KIM-1 staining in kidneys from SARS-CoV-2 infected k18hACE2 mice that received ACE2 618-DDC-ABD pre- and post-viral inoculation.

- **Table 1.** Scoring system in BSL-3 facility for health evaluation of mice infected with SARS-CoV-2.

**Figure S1.** Examples of GFAP and IBA1 staining in the brain of SARS-CoV-2 inoculated k18hACE2 mice.

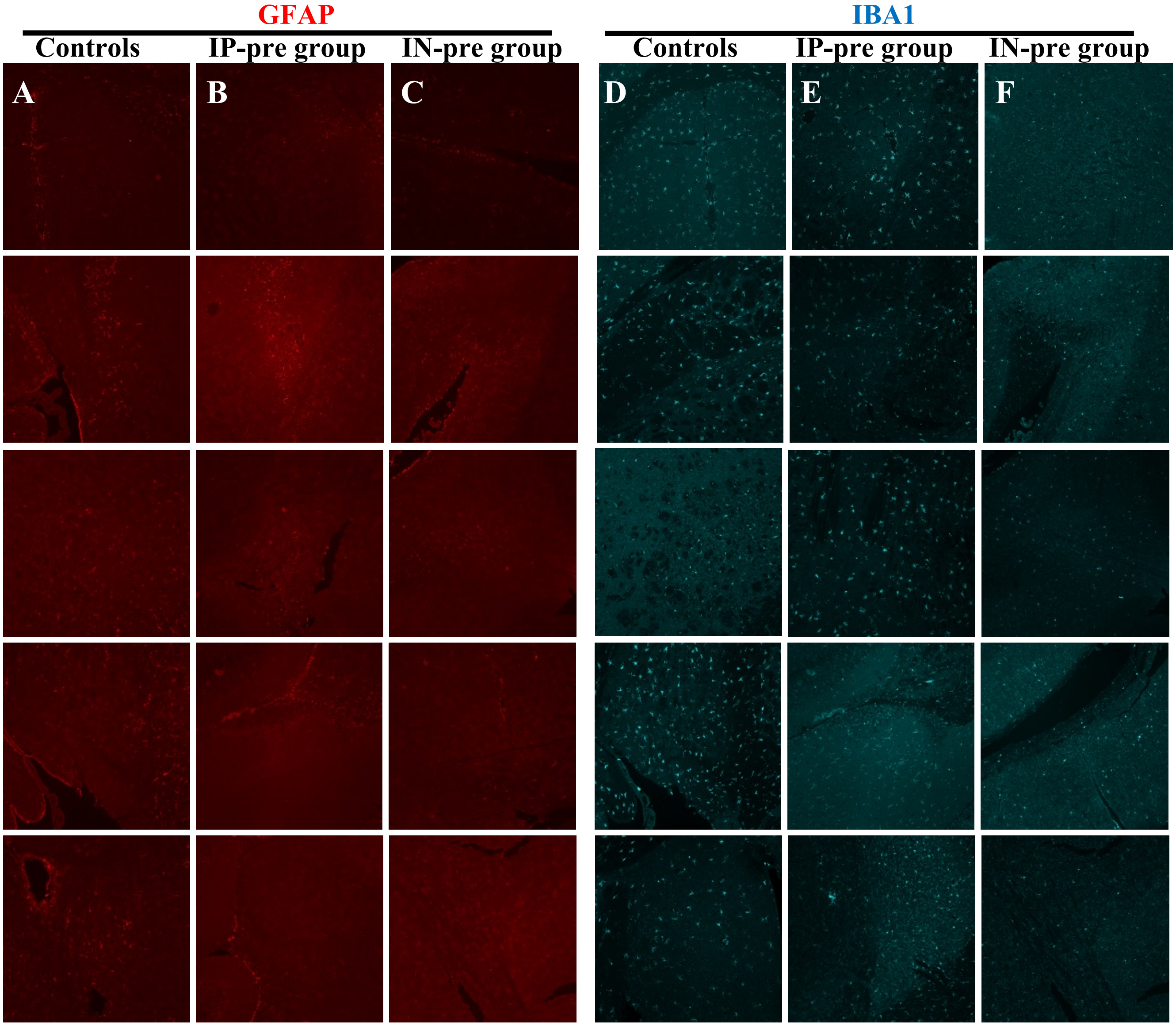


**Figure S1.**
**Panel A-C.** GFAP staining (red) is strong in the brain of 5 infected, untreated controls (**A**), whereas it is partially reduced in the brain of 5 mice from the IP-pre group that received ACE2 618-DDC-ABD (**B**). In 5 mice from the IN-pre group, GFAP staining is further reduced or almost completely absent in some mice (**C**).
**Panel D-F.** IBA1 staining (blue) is strong in untreated, infected controls (**D**) and mice from the IP-pre group that received ACE2 618-DDC-ABD (**E**). In mice from the IN-pre group, by contrast, GFAP staining is reduced (**F**). All photomicrographs were taken at 20x magnification, please magnify to see the differences better.

**Figure S2.** Representative brain histopathology of SARS-CoV-2 infected k18hACE2 mice that received ACE2 618-DDC-ABD only post viral inoculation.

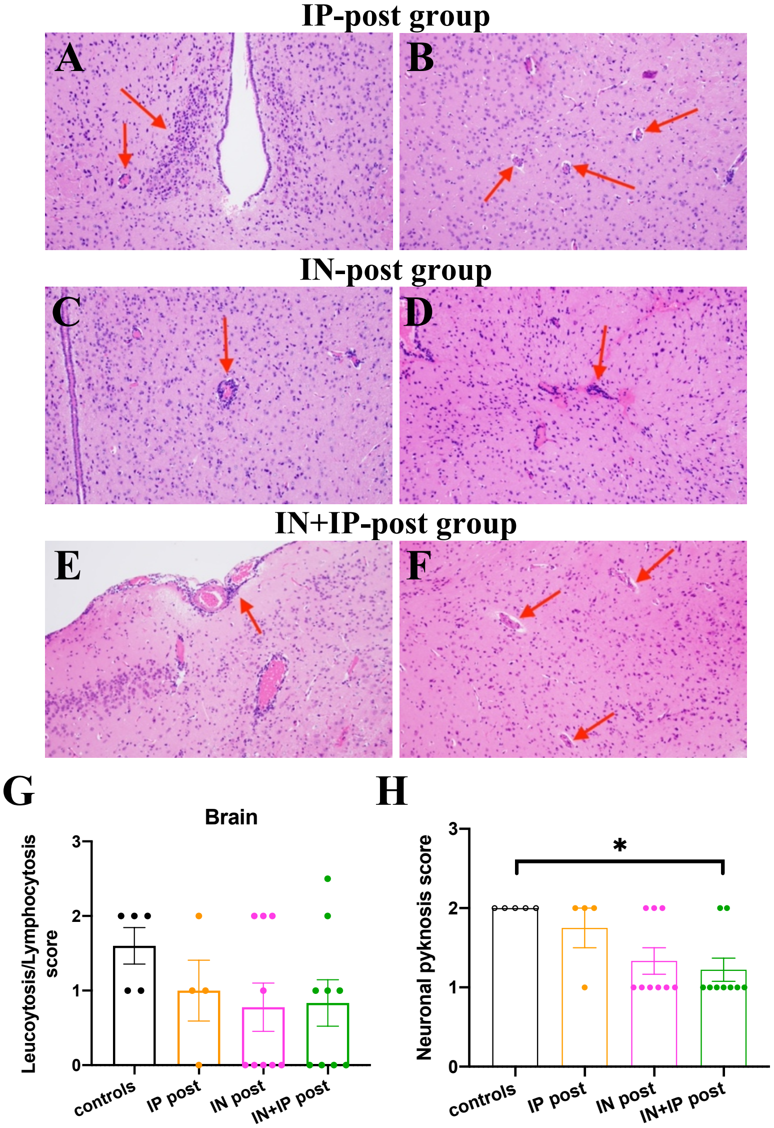


**Figure S2.**

**Panel A, B.** In mice from the IP-post group that received ACE2 618-DDC-ABD, perivascular and parenchymal inflammation (**A**) and endothelial hypertrophy (**B**) were seen in the hypothalamus.

**Panel C, D.** In mice from the IN-post group, perivascular lymphocytosis was seen in the lateral hypothalamus (**C**) and lateral cortex (**D**).

**Panel E, F.** In mice from the IN+IP-post group, perivascular and leptomeningeal lymphocytosis in the lateral cortex (**E**) and endothelial hypertrophy in the lateral hypothalamus (**F**) were seen. All photomicrographs were taken at 20x magnification.

**Panel G.** The histopathological score for leukocytosis/lymphocytosis was the highest in infected untreated controls (black). It was decreased in all treated groups (color), but these differences were not statistically significant.

**Panel H.** The histopathological score for neuronal pyknosis was highest in infected untreated controls (black). It was lower in the IP-post group (orange), the IN-post (pink) and IN+IP-post group (green). The differences between the groups were statistically significant for the IN+IP-post group (green) as compared to controls ($p=0.0370$). Mean ± SEM are shown. Significance was calculated by one-way ANOVA followed by Dunn's multiple comparisons test.

**Figure S3.** Representative lung histopathology in SARS-CoV-2 infected k18hACE2 mice that received ACE2 618-DDC-ABD only post viral inoculation.

**IP-post group**

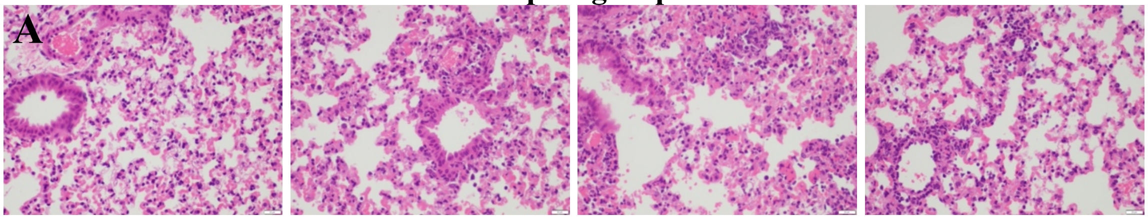


**IN-post group**

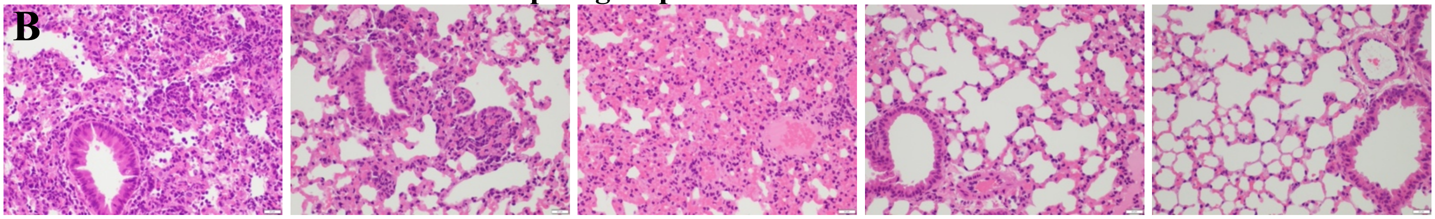


**IN+IP-post group**

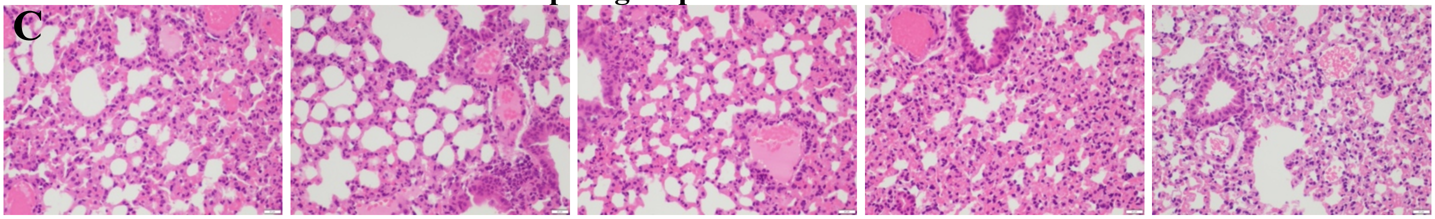


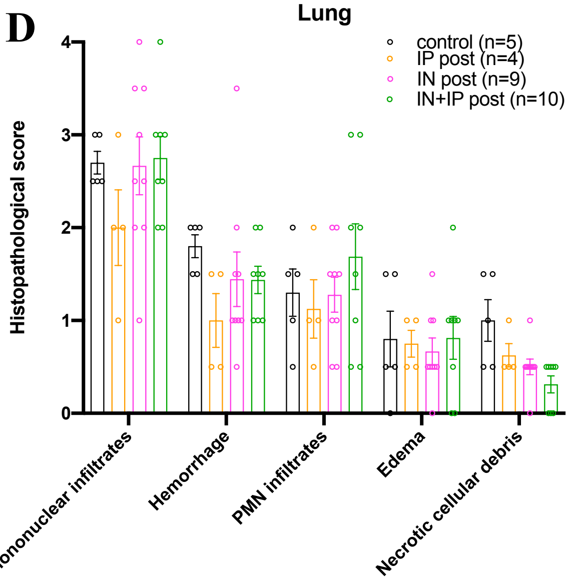


**Figure S3.**

**Panel C.** Four different mice from the group that received **ACE2 618-DDC-ABD post viral inoculation via IP** show marked perivascular mononuclear infiltrates, alveolar hemorrhage and scattered neutrophils and resemble mostly the infected untreated controls.

**Panel D.** Five different mice from the group that received **ACE2 618-DDC-ABD post viral inoculation via IN** also show marked perivascular mononuclear infiltrates, alveolar hemorrhage and scattered neutrophils. In some mice from this group, however, lung histopathology was improved as compared to the infected untreated controls.

**Panel E.** Five different mice from the group that **received ACE2 618-DDC-ABD post viral inoculation via IN+IP** show marked perivascular mononuclear infiltrates, alveolar hemorrhage and scattered neutrophils and are similar to the infected untreated controls. All photomicrographs were taken from H&E-stained sections at 40x magnification, scale bar = 500um.

**Panel F.** The lung histopathology scores for mononuclear infiltrates, edema and PMN infiltrates and edema in the infected untreated controls (black) are not different as compared to the post-treated mice with ACE2 618-

DDC-ABD (color). The scores for hemorrhage necrotic cellular debris are lower in the treated groups (color) as compared to controls (black). These differences were not significant. Mean ± SEM are shown. Significance was calculated by mixed-effects analysis followed by Tukey's multiple comparisons test. Organs from one mouse each in the IP and IN group could not be obtained.

**Figure S4.** Representative kidney histopathology of SARS-CoV-2 infected k18hACE2 mice.

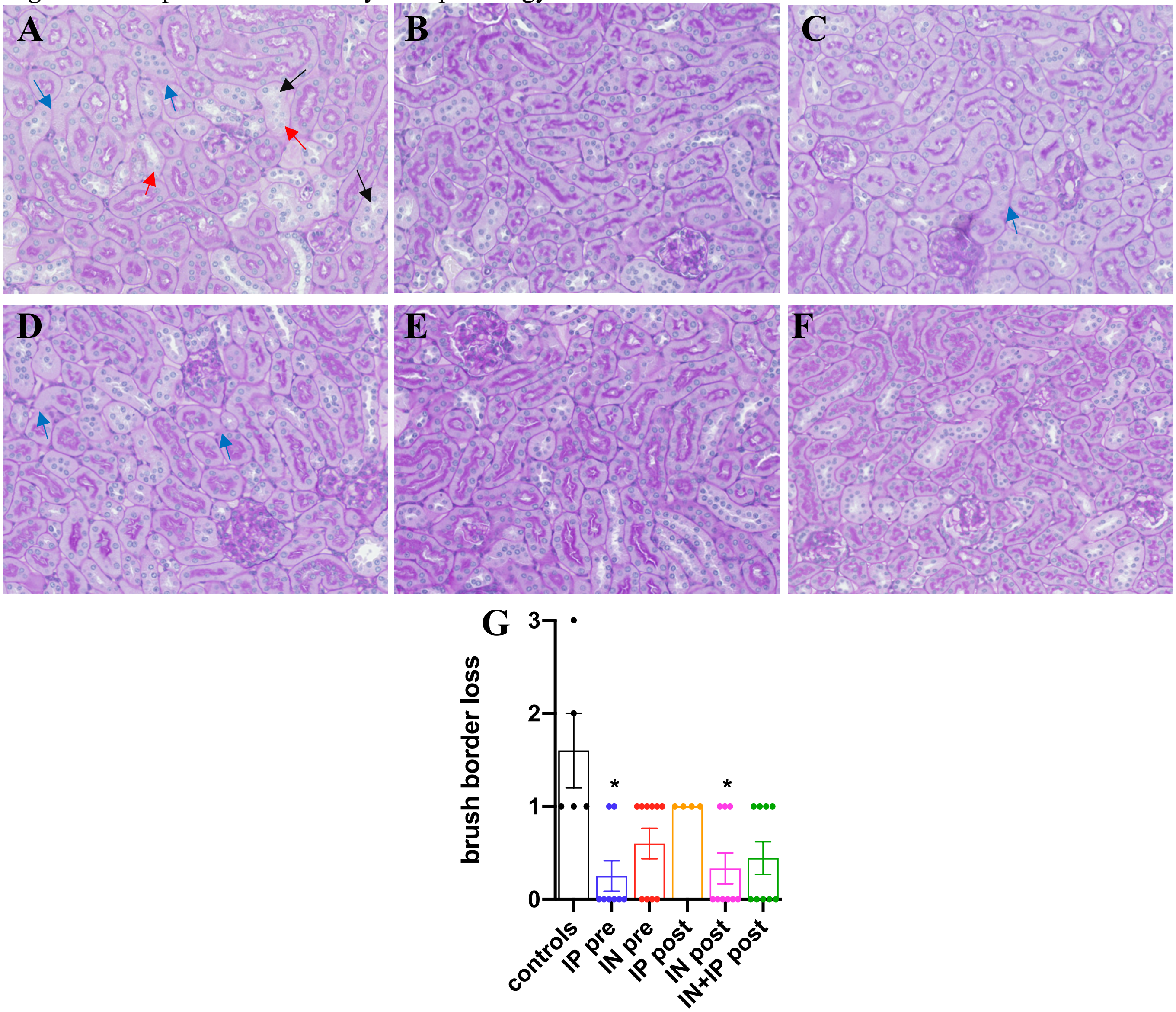


**Figure S4.**

**Panel A.** Example of a PAS-staining of the kidney from a SARS-CoV-2 infected, untreated mouse showing focal attenuation of proximal tubular brush border (black arrow), cytolysis (red arrow) and mild basement membrane disruption (blue arrow).

**Panel B, C.** Example of PAS-staining of kidneys from infected mice that received ACE2 618-DDC-ABD pre- and post-viral inoculation either intraperitoneally (**B**) or intranasally (**C**) showing almost intact proximal tubules.

**Panel D-F.** Example of PAS-staining of kidneys from infected mice that received ACE2 618-DDC-ABD only post-viral inoculation either intraperitoneally (**D**), intranasally (**E**) or intranasally + intraperitoneally (**F**) showing near normal proximal tubules. 20x magnification.

**Panel G.** The score for brush border loss is highest in infected untreated controls (black) and reduced in the groups that received ACE2 618-DDC-ABD (color). The difference reaches statistical significance only for controls as compared to the IP-pre group (blue, $p=0.02$) and the IN-post group (pink, $p=0.04$). Mean ± SEM are shown. Significance was calculated by one-way ANOVA followed by Dunn's multiple comparisons test.

**Figure S5.** NGAL staining in kidneys from SARS-CoV-2 infected k18hACE2 mice that received ACE2 618-DDC-ABD pre- and post-viral inoculation.

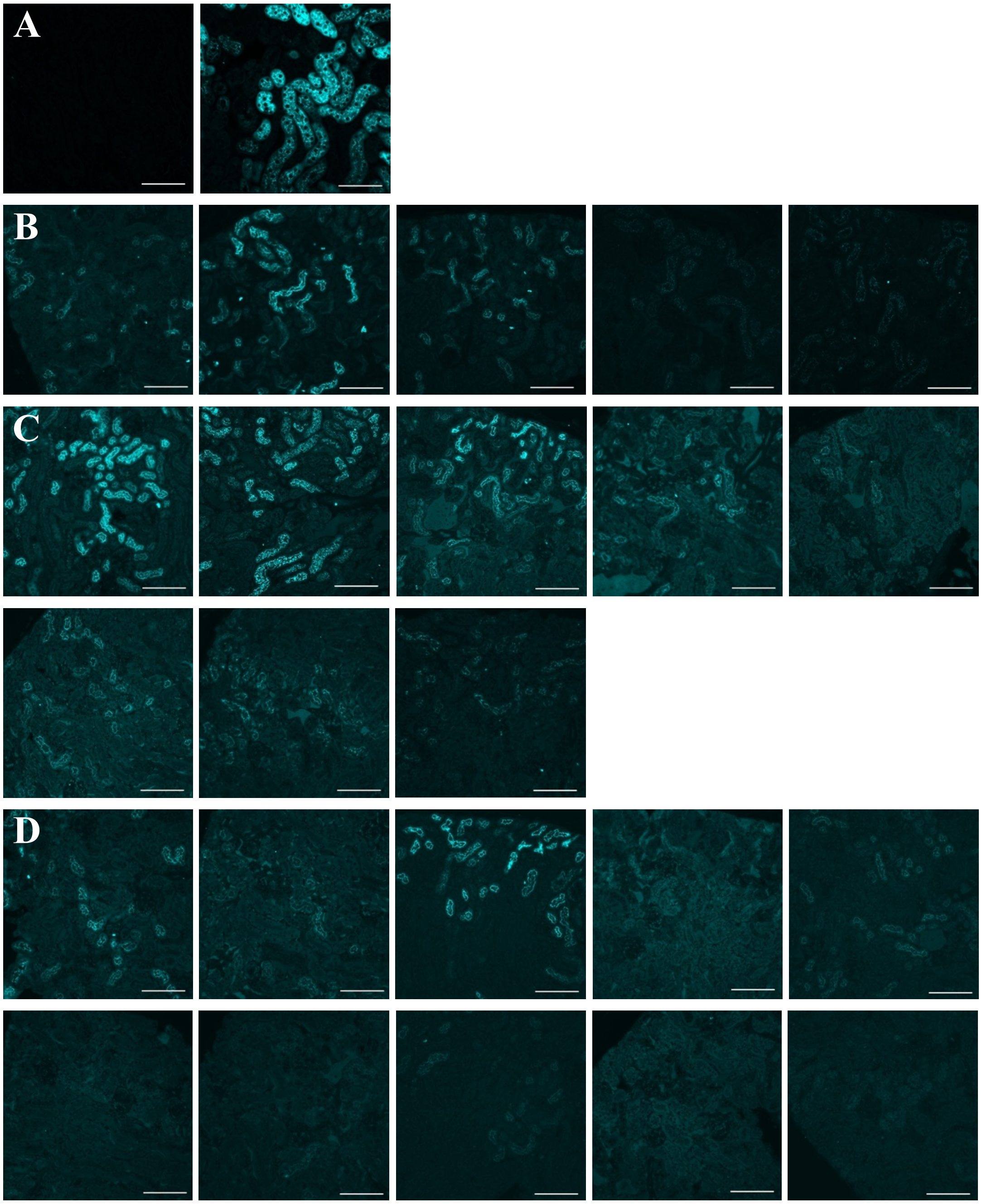


**Figure S5.**
**Panel A.** Examples of NGAL staining in a normal kidney (absent, **left**) and in a kidney with AKI 48 hours after ischemia-reperfusion injury (30 minutes) (strong, **middle**).
**Panel B.** In kidneys of k18hACE2 mice **infected and untreated** (n=5), NGAL staining was clearly evident in 4 mice and weak in the last one.
**Panel C.** In kidneys of mice that received ACE2 618-DDC-ABD prior to viral inoculation via IP (**IP-pre group**, n=8, organs of two mice could not be obtained), NGAL staining was evident in 3 mice and weak in the 5 others.
**Panel D.** In kidneys of mice that received ACE2 618-DDC-ABD prior to viral inoculation via IN (**IN-pre group**, n=10), NGAL staining was evident in 3 mice, and weak or undetectable in the remaining 7 mice.
40x magnified, scale bar = 100um. Please zoom in to see the differences better.

**Figure S6.** NGAL staining in kidneys from SARS-CoV-2 infected k18hACE2 mice that received ACE2 618-DDC-ABD only post viral inoculation.

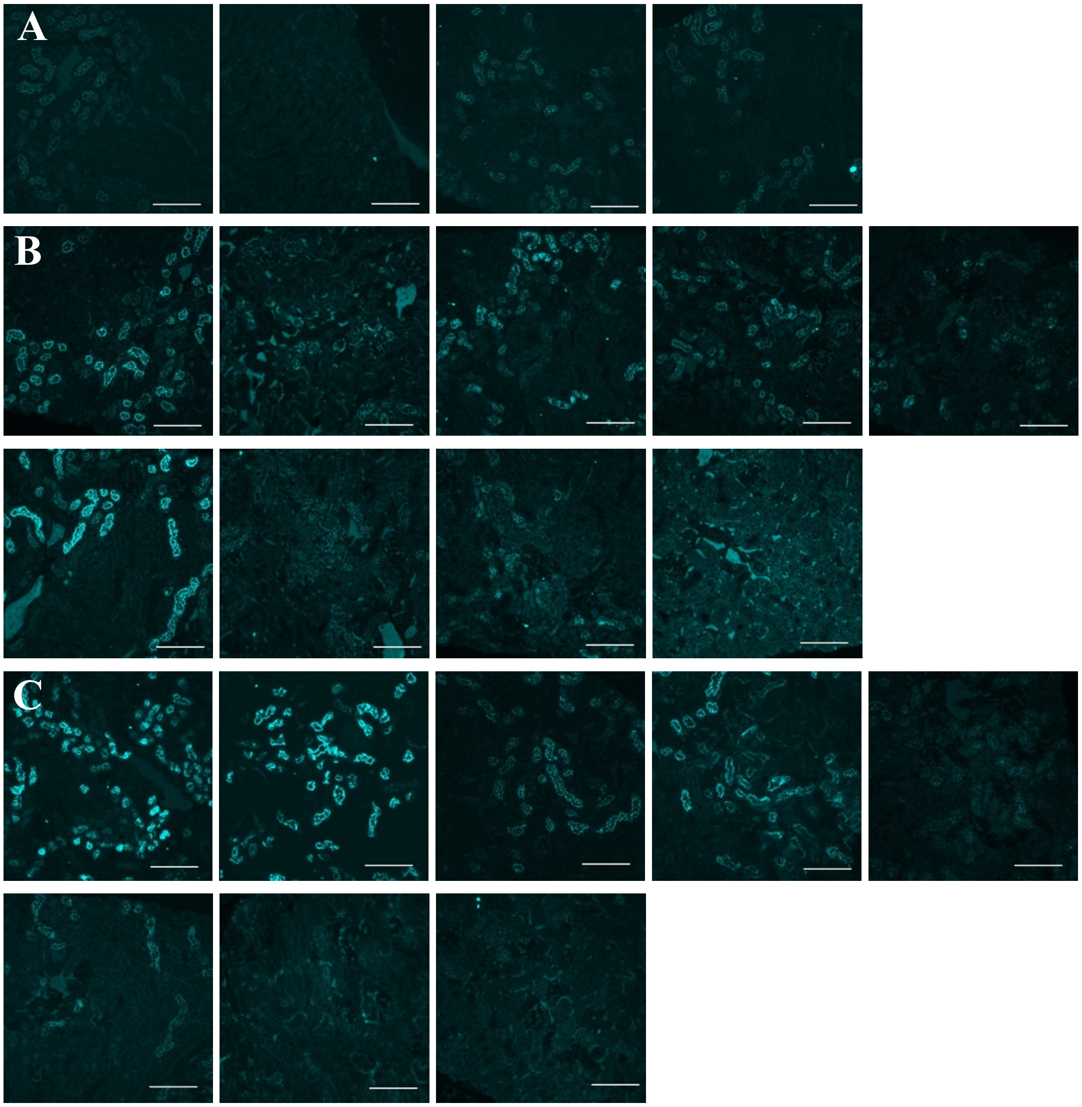


**Figure S6.**
**Panel A.** In mice that received ACE2 618-DDC-ABD post viral inoculation via IP (**IP-post group**, n=4), NGAL staining was present but reduced as compared to controls.
**Panel B.** In mice that received ACE2 618-DDC-ABD post viral inoculation via IN (**IN-post group**, n=9), NGAL staining was present in most mice, whereas some mice from this group had reduced or undetectable NGAL staining.
**Panel C.** In mice that received ACE2 618-DDC-ABD post viral inoculation via IN+IP (**IN+IP-post group**, n=8), NGAL staining was present in most mice, whereas some mice from this group had reduced or undetectable NGAL staining.
40x magnified, scale bar = 100um. Organs from one mouse per group could not be obtained, and kidney tissue quality was not suitable for staining in one mouse from the IN+IP-post group. Please zoom in to see the differences better.

**Figure S7.** KIM-1 staining in kidneys from SARS-CoV-2 infected k18hACE2 mice that received ACE2 618-DDC-ABD pre- and post-viral inoculation.

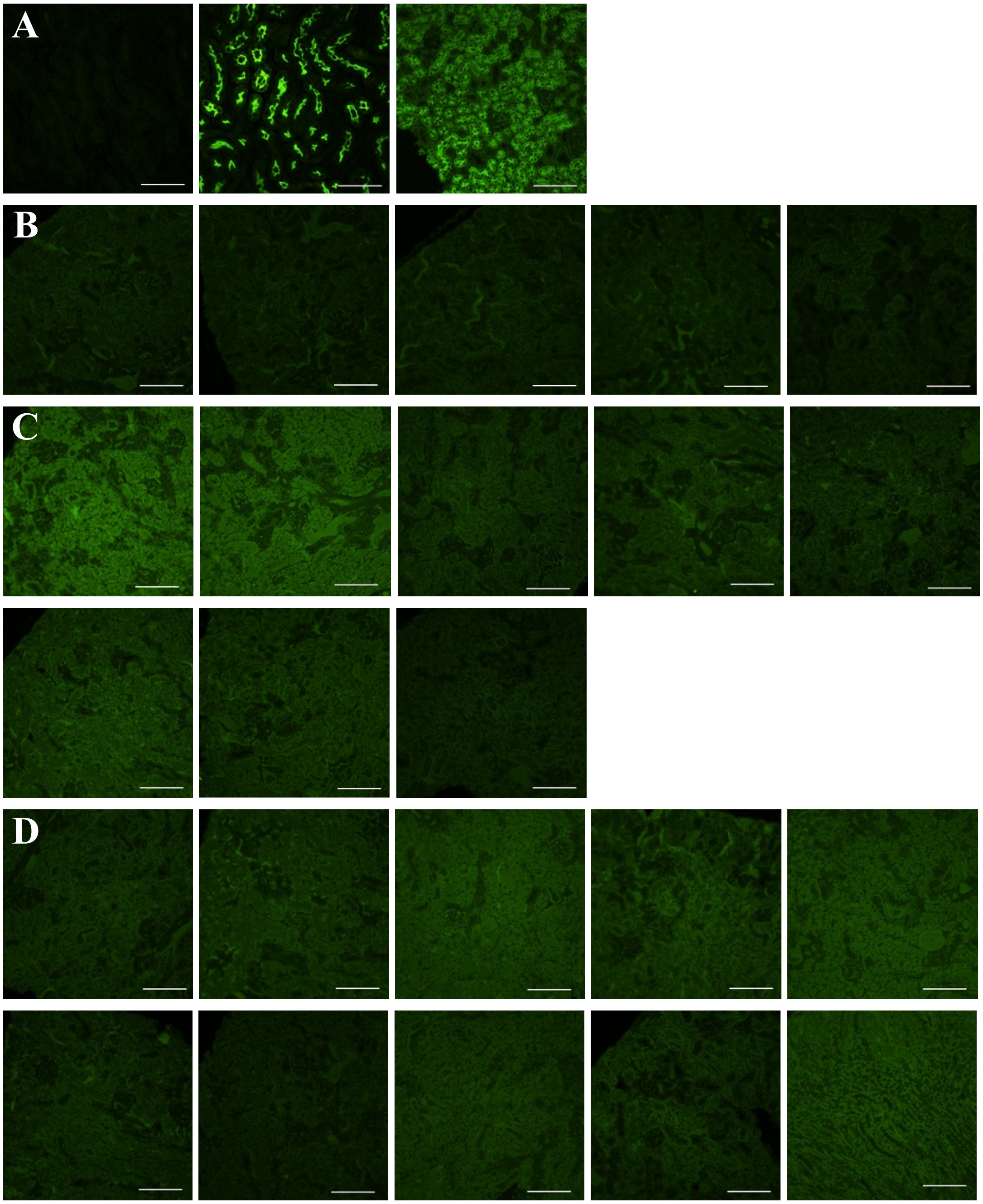


**Figure S7.**

**Panel A.** Examples of KIM-1 staining in a normal kidney (absent, **left**), in a kidney with AKI 48 hours after ischemia-reperfusion injury (30 minutes) (strong, **middle**) and a kidney of a k18hACE2 mouse infected with SARS-CoV-2 (less strong, **right**).

**Panel B.** In the kidney of SARS-CoV-2 **infected untreated** k18hACE2 mice (n=5), KIM-1 staining was absent.

**Panel C, D.** In the treated groups that received ACE2 618-DDC-ABD pre- and post-viral inoculation (**IP-pre and IN-pre group**, n=8 and n=10, respectively), KIM-1 staining was absent.

40x magnified, scale bar = 100um. Organs from two mice in the IP-pre group could not be obtained. Please zoom in to see the differences better.

**Table 1.** Scoring system in BSL-3 facility for health evaluation of mice infected with SARS-CoV-2.

| Score | Description |
|---|---|
| **0** | Pre-inoculation: mice are bright, alert, active, normal fur coat and posture |
| **1** | Post-inoculation (PI): mice are bright, alert, active, normal fur coat and posture, no weight loss |
| **1.5** | Mice present with slightly ruffled fur but are active, or weight loss might occur but <2.5%, recovery can be expected |
| **2** | Ruffled fur or less active or weight loss <5%, recovery might occur |
| **2.5** | Ruffled fur or not active but moves when touched or hunched posture or difficulty breathing or weight loss 5-10%, Recovery is unlikely but still might occur |
| **3** | Ruffled fur or inactive but moves when touched or difficulty breathing or weight loss at 11-20%, recovery is not expected |
| **4** | Ruffled fur or positioned on its side or back or dehydrated or difficulty breathing or weight loss >20% or labored breathing, recovery is not expected |
| **5** | death |
